# DNA-based nanocarriers to enhance the optoacoustic contrast of tumors *in vivo*

**DOI:** 10.1101/2020.08.07.235705

**Authors:** James Joseph, Kevin N. Baumann, Alejandro Postigo, Laura Bollepalli, Sarah E. Bohndiek, Silvia Hernández-Ainsa

## Abstract

Optoacoustic tomography (OT) enables non-invasive deep tissue imaging of optical contrast at high spatio-temporal resolution. The applications of OT in cancer imaging often rely on the use of molecular imaging contrast agents based on near infrared dyes to enhance contrast at the tumor site. While these agents afford excellent biocompatibility and minimal toxicity, they present limited optoacoustic signal generation capability and rapid renal clearance, which can impede their tumor imaging efficacy. In this work, a synthetic strategy to overcome these limitations utilizing biodegradable DNA-based nanocarrier (DNA-NC) platforms is introduced. DNA-NCs enable the incorporation of near infrared dyes (in this case, IRDye 800CW) at precise positions to enable fluorescence quenching and maximize optoacoustic signal generation. Furthermore, these DNA-NCs show a prolonged blood circulation compared to the native fluorophores, facilitating tumor accumulation by the Enhanced Permeability and Retention (EPR) effect. *In vivo* imaging of tumor xenografts in mice following intravenous administration of DNA-NCs revealed enhanced OT signals at 24h when compared to free fluorophores, indicating promise for this method to enhance the optoacoustic signal generation capability and tumor uptake of clinically relevant near infrared dyes.

## 1. Introduction

Optoacoustic tomography (OT) combines the rich contrast of optical imaging with highspatial resolution and penetration depth of ultrasound.^[1, 2]^ Upon absorption of an irradiating light pulse, transient thermoelastic expansion gives rise to an acoustic wave that can be detected at the surface of the imaged tissue. Acquiring OT data at multiple wavelengths enables the visualization and quantification of anatomical and functional information from endogenous tissue chromophores, such as hemoglobin, lipids and melanin, as well as offering molecular imaging capabilities using exogenous contrast agents.^[3-8]^ For molecular imaging, particularly in clinical applications, OT often utilizes near infrared (NIR) small molecule dyes having excellent biocompatibility and minimal toxicity, which can enhance the contrast for visualization of diseases such as cancer.^[9]^ Unfortunately, such small molecule dyes present limited optoacoustic signal generation capabilities and rapid clearance from the systemic circulation ^[10]^, affording limited opportunity for accumulation at the target imaging site.^[11, 12]^

Advances in nanomedicine have led to increasing innovation in a range of different types of biocompatible nanocarriers (NCs) for disease diagnosis and therapy,^[13-16]^ such as liposomes or polymeric nanoparticles. These nanocarriers are capable of packaging small molecule dyes, which can enhance their optical absorption properties for OT.^[17, 18]^ The integration of such small molecule dyes into nanocarriers presents the further advantage of increasing the overall contrast agent size, which helps to lower their renal elimination rate,^[10]^ hence prolonging blood half-life and promoting their delivery to the tumor site by the Enhanced Permeability and Retention (EPR) effect.^[19, 20]^

Structural DNA nanotechnology renders a promising strategy to engineer biocompatible NCs with customized structures and sizes in a highly reproducible fashion.^[21]^ DNA-based NCs have demonstrated significant advantages over other engineered nanoparticles for the transport and delivery of various diagnostic and therapeutic agents for *in vivo* applications in cancer.^[22-27]^ In the context of *in vivo* cancer imaging, DNA-NCs have been loaded with a variety of contrast agents for different modalities, including dyes for fluorescence imaging,^[28]^ radioisotopes for SPECT^[29]^ and PET,^[30]^ as well as gold nanoparticles for OT.^[23]^

Based on the knowledge that distance-dependent quenching of small molecule NIR dyes yields significant enhancement of OT signals,^[31, 32]^ we hypothesized that DNA-NCs could be used to construct biodegradable OT contrast agents with high optoacoustic signal generation and tumor uptake properties. Using precise positioning of IRDye 800CW^[33]^ fluorophore molecules at very close proximities within DNA nanostructures, we obtained a substantial OA signal enhancement and tailored the effective size of these nanostructures to offer prolonged blood clearance times compared to the free dyes, thereby enhancing tumor uptake. We performed *in vivo* OT in mice bearing tumor xenografts to demonstrate modulation of renal clearance rate and enhanced tumor to background ratio for the DNA-NCs.

## 2. Results

### 2.1. DNA-NCs design, preparation and structural characterization

We first adapted two DNA-NCs, namely NC8 and NC6 (**Figure 1a**), from a previously reported design^[34]^ following the single-stranded tile method.^[35]^ The NCs contain 8 and 6 DNA oligonucleotides respectively, with NC8 presenting larger structural rigidity and size compared to NC6 (see Table S1 in Supporting Information). Optical absorption capability was incorporated by assembling the structures using four oligonucleotides functionalized with the IRDye 800CW fluorophore at their terminal sides. Two distinct classes of NC8 and NC6 were synthesized (Q+ and Q-) having a different number of dyes and different inter-dye separation distances to confirm the quenching-based origin of the OT signal enhancement. Specifically, NC8Q+ and NC6Q+ contain four fluorophores arranged in pairs with minimal inter-dye separation distance to enable H-dimer formation,^[36-38]^ whereas NC8Q- and NC6Q- contain only two fluorophores with larger inter-dye separation distance (Figure 1a). The NCs were assembled following thermal gradients used in previous reports ^[39]^ (see Experimental Section). Polyacrylamide gel electrophoresis (PAGE) and Dynamic Light Scattering (DLS) were used to assess the correct folding of NC6 and NC8. On the one hand, PAGE shows a distinct band for each NC, with NC8 (Q+ and Q-) running slower than NC6 (Q+ and Q-) due to the larger molecular weight and size (Figure 1b). DLS data (Figure 1c) show different peak sizes for the NCs, given by their hydrodynamic diameters. By Gaussian-fitting the size distributions, the average hydrodynamic diameter (in number) was determined as 9.5 nm ± 2.3 nm and 6.6 nm± 1.5 nm for NC8 and NC6 respectively. Prior to their optical characterization, the stability of NCs incubated up to 48 hours in cell culture media containing 10% fetal calf serum (FCS) was tested to assess their suitability for further *in vivo* imaging studies. NC8Q+ showed remarkably lower stability compared NC6Q+ (Figure 1d).

**Figure 1.**
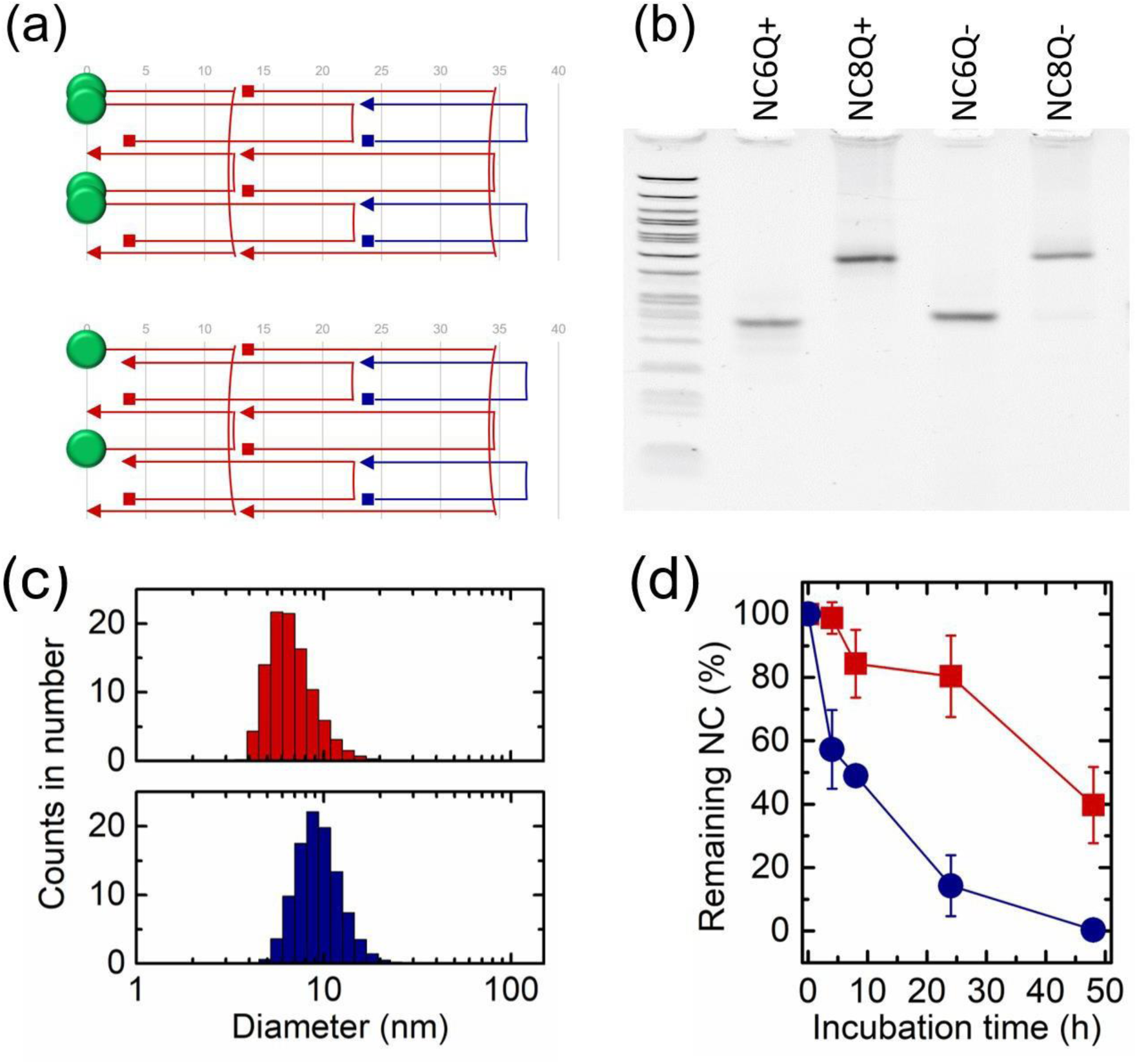
DNA NC design and fabrication. (a) Graphical presentation of the layout of the DNA nanocarriers (NCs). NC6 is obtained by depleting the two oligos represented in blue color. IRDye 800CWfluorophores are shown as green disks. Top panel and bottom panel gather the Q+ or Q-NC respectively. (b) PAGE of the different NCs. A 50 bp ladder is included in the first lane. (c) Hydrodynamic diameter (in number) obtained by DLS of the NC6 (red graph) and NC8 (blue graph). (d) Remaining NC6 (red data) and NC8 (blue data) estimated by PAGE analysis after incubation with DMEM containing 10% FCS at different time points.

### 2.2. Optical characterisation of DNA-NCs reveals high quenching efficiencies

Given the enhanced stability observed in the NC6 structures, we decided to proceed to further optical, cellular and *in vivo* studies using smaller structures. In addition to NC6, we also prepared a simpler construct consisting of 21-base pairs long duplex (NC2); NC2Q+ bearstwo IRDye 800CW fluorophores both arranged at one side whereas NC2Q- was only labelled with one fluorophore (see Table S1 in Supporting Information). The optical response of the NCs was initially evaluated using optical absorption and fluorescence spectroscopy. Optical absorption spectra of NC6Q- (light red line in **Figure 2a**) showed the characteristic absorption peak (778 nm) of the IRDye 800CW. In the case of NC6Q+ (dark red line in Figure 2a) two peaks (maxima at 705 nm and 778 nm) were observed, supporting the formation of H-dimers due to the close positioning of NIR dye molecules. On the other hand, fluorescence emission was quenched from NC6Q- (light red spectrum) to NC6Q+ (dark red spectrum) (Figure 2b). The same behavior was found for the absorbance and emission spectra of NC2Q+ and NC2Q- (see Figure S1a,b in Supporting Information). Fluorescence quenching efficiencies of 80 ± 4% and 86 ± 2% were observed for NC6 and NC2 respectively. These values were determined by comparing the intensities at the maximum of the fluorescence spectra of the NCQ+ and NCQ-forms at matching dye concentrations recorded upon excitation at 778 nm (see Experimental Section).

**Figure 2.**
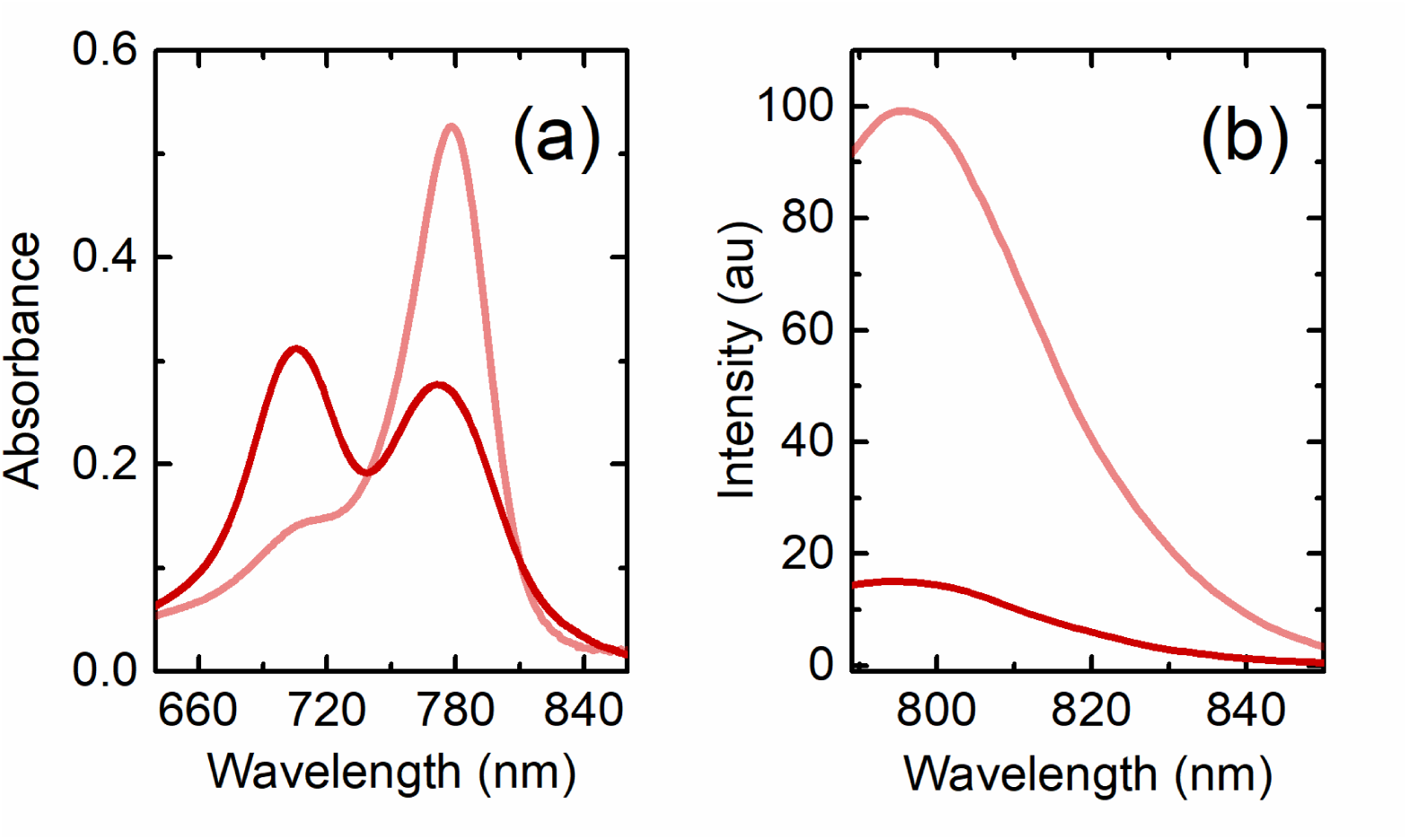
Optical characterization of the DNA NCs. (a) Absorbance and (b) emission spectra of NC6Q+ (in dark red) and NC6Q- (in light red).

### 2.3. Phantom studies using OT reveal optoacoustic signal enhancement

The OT response of the NCs at depths relevant for further *in vivo* studies was initially evaluated using tissue mimicking phantoms. Samples were encapsulated into agar-based tissue mimicking phantoms with defined optical properties (scattering coefficient of 5cm^-1^ and absorption coefficient of 0.05cm^-1^ at 700 nm) as previously described^[31, 32]^ (see Experimental Section). The OT spectra of NCQ- and NCQ+ structures (**Figure 3a** for NC6 and supporting information Figure S1c for NC2) showed the expected signal peaks in accordance with the measured absorbance (Figure 2a for NC6 and Figure S1a in Supporting Information for NC2). The area under the curve (AUC), which gives the integrated OT mean pixel intensities, showed an average 1.6-fold OA enhancement (p=0.023) for NC6Q+ when compared to NC6Q- (Figure 3b) and 2.1-fold OA enhancement (p=0.013) for NC2Q+ when compared to NC2Q- (Figure 3c).

**Figure 3.**
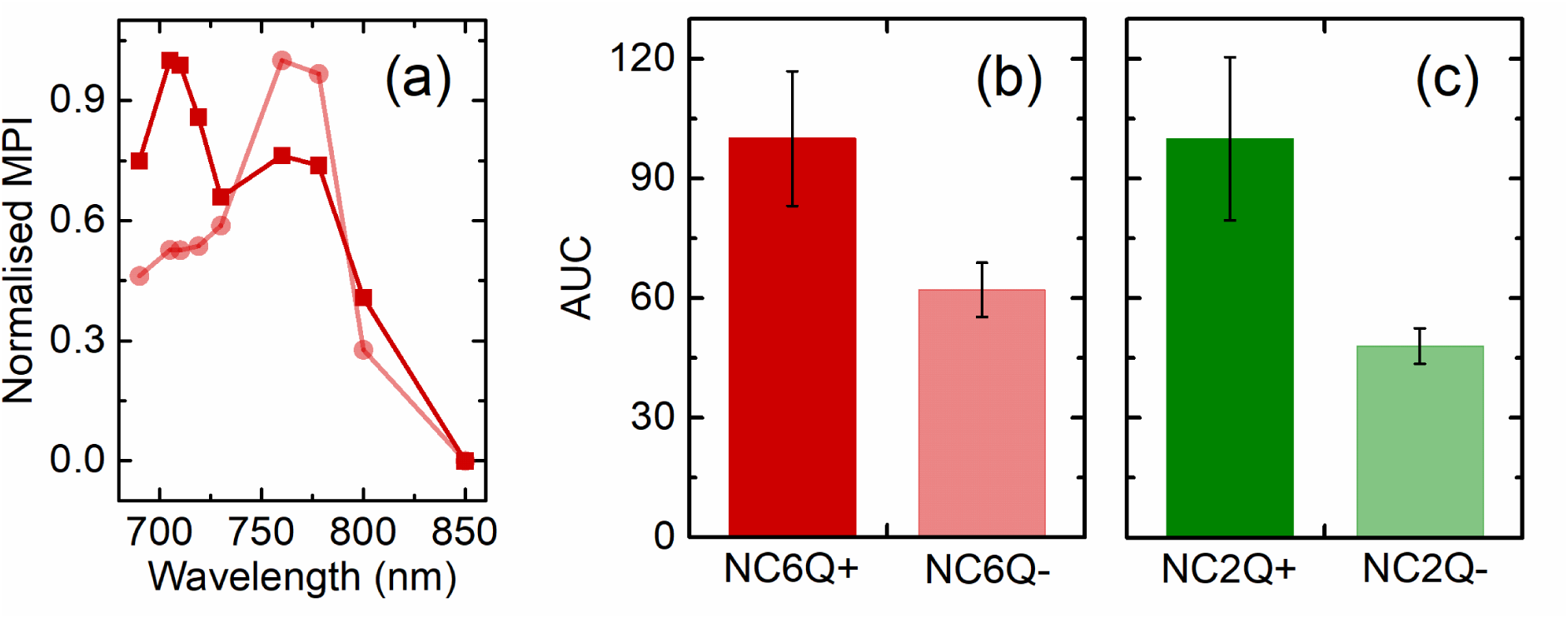
(a) Phantoms OT spectra of NC6Q+ (in dark red) and NC6Q- (in light red). (b) AUC data obtained from the OT spectra of (b) NC6 and (c) NC2 samples. AUC data correspond to the average and standard deviation of 3 different samples. MPI: mean pixel intensity.

### 2.4. DNA-NCs did not influence cellular proliferation

In order to evaluate the potential impact of the NCs in the tumor bed *in vivo*, NCs were incubated with the MDA-MB-231 cell line for a proliferation test using a real-time cell health and viability analyzer (see Experimental Section). Cell confluence was measured for up to 168 hours. Different concentrations of DNA-NCs were assessed, with the highest concentrations designed to exceed those that would be expected to be present locally within tissue during the subsequent *in vivo* studies. As observed in **Figure 4**, no significant differences were found between the confluence rate of the cells incubated with either of the NCs and the control in PBS.

**Figure 4.**
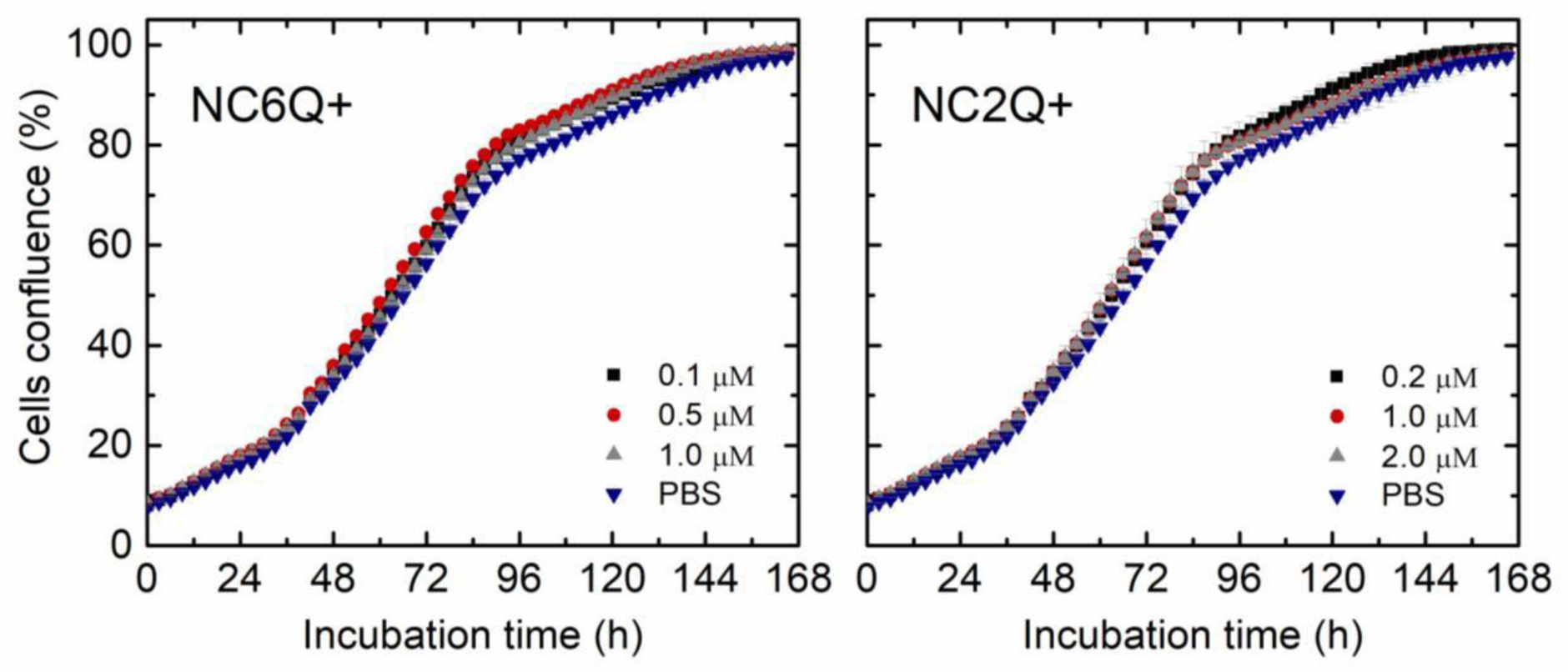
Confluence (%) at different incubation times of breast cancer MDA-MB-231 cells in the presence of different concentrations of NC6Q+ and NC2Q+. Data and error bars correspond to the average and standard deviation of 4 measurements.

### 2.5. OT reveals extended renal clearance time of DNA-NCs compared to free dye

To investigate whether the incorporation of the dye into the NCs could increase its *in vivo* blood retention time,^[40]^ we investigated their biodistribution kinetics by performing fast multispectral optoacoustic tomography of healthy mice administered with the free fluorophore IRDye 800CW, NC2Q+ and NC6Q+. *In vivo* injected concentrations were 32 μM for the free dye, 16 μM for NC2Q+ and 8 μM NC6Q+, to give equivalent dye concentration in the intravenous administration injection into the nude (BALB/c nu/nu) mice (n=3 for each group). Fast OT cross-sectional images were acquired at multiple wavelengths along the mouse body. The clearance of the probes was found to occur primarily in the kidneys (**Figure 5a)** and liver (Figure 5b) though both DNA-NCs and free dye primarily exhibited a renal clearance pathway, in agreement with the studies reported earlier for the IRDye 800CW fluorophore.^[41]^

**Figure 5.**
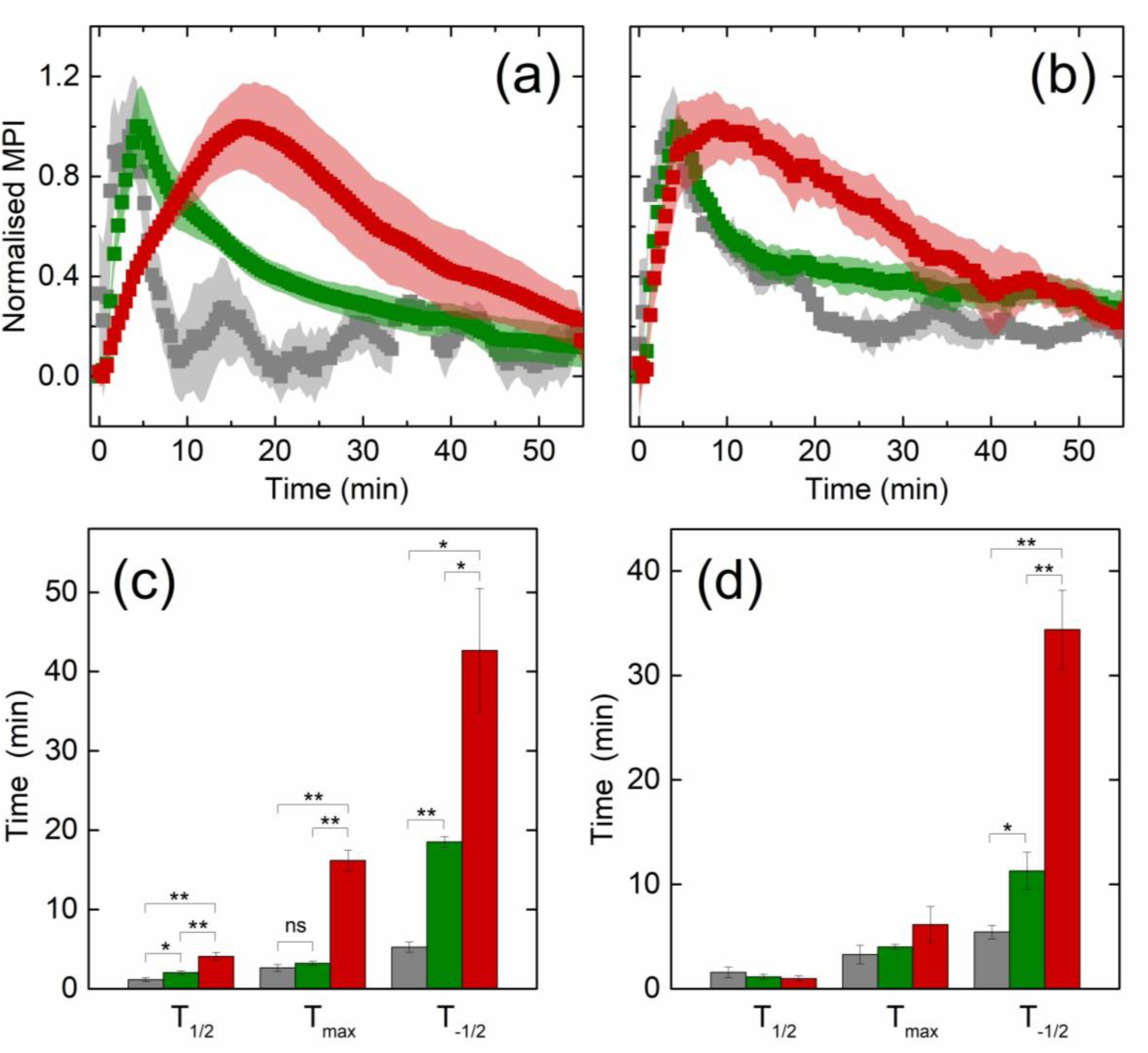
Temporal evolution of optoacoustic signal from kidney and liver obtained from fast MSOT studies. Normalized mean pixel intensity (MPI) values estimated from (a) renal cortex and (b) liver cross-sections and time resolved OT signal (in minutes) obtained from the injected probes in (c) renal cortex and (d) liver cross-sections. Red data correspond to NC6Q+, green data correspond to NC2Q+ and grey data correspond to IRDye 800CW. Data correspond to the average and SEM values obtained from 3 mice. Extracted metrics included: time to half maximum (T_1/2_); Time to maximum (T_max_); Time from maximum to half maximum (T-_1/2_). n=3

In order to determine the rate of this clearance process, the evolution of the OT signal over time from the renal cortex and liver was assessed. The quantification of the signal dynamics from Figure 5a and 5b for the renal cortex and liver gave the kinetic parameters presented in Figure 5c and 5d, respectively (see also Table S2 in Supporting Information). As expected IRDye 800CW rapidly clears via kidneys, whereas NC2Q+ and NC6Q+ exhibit a more complex clearance pathway and extended clearance time. Uptake in the renal cortex increases rapidly immediate after injection for IRDye 800CW, also decaying rapidly thereafter. Uptake for NC2Q+ and NC6Q+ evolves over respectively longer time periods and decays more slowly, indicating a prolonged retention, especially for NC6Q+ (Figure 5a). In liver, IRDye 800CW, NC2Q+ and NC6Q+ show a similar rate in their uptake (Figure 5b). However, in contrast to IRDye 800CW and NC2Q+, NC6Q+ renders decelerated signal decay, indicating a slower clearance rate from the liver. Further, time resolved OT signal from kidneys show significant differences in the uptake and clearance of NC6Q+ when compared to IRDye 800CW and NC2Q+ at defined time points. However, differences in the uptake and clearance of the probes from liver at T_1/2_ and T_max_ were found not to be significant.

### 2.6. OT of tumor bearing mice demonstrates enhanced tumor uptake

Based on the extended blood half-life of NC6Q+, this DNA-NC was compared to free IRDye 800CW in tumor-bearing mice. For *in vivo* tumor imaging studies, NC6Q+ and IRDye 800CW at the same dye concentrations were intravenously administered via tail vein of female nude (BALB/c nu/nu) mice bearing subcutaneous MDA-MB-231 tumors (n=3 for each group). Prior to the administration of the contrast agents, baseline measurements for both OT and fluorescence imaging were performed. OT was performed during the administration of the contrast agent and up to 20 minutes post injection, immediately followed up with fluorescence imaging. OT and fluorescence imaging were also carried out at 24h and 48h time points. The OT signals for NC6Q+ and IRDye 800CW remained evident within the tumor bed at later time points **(Figure 6a)**, which may be due to the EPR effect. NC6Q+ exhibits OT mean pixel intensities 2.7-fold larger than the free dye at 24h (p=0.045) and remains elevated at 48h (Figure 6b). The *in vivo* fluorescence imaging capabilities of the NCs were also assessed (data not shown), where NC6Q+ structures are expected to exhibit low fluorescence due to the self-quenching of the IRDye 800CW fluorophores. Fluorescence intensities for NC6Q+ and free IRDye 800CW showed similar values at 24 and 48 hours post injection, quantified from the surface weighted fluorescence signals.

**Figure 6.**
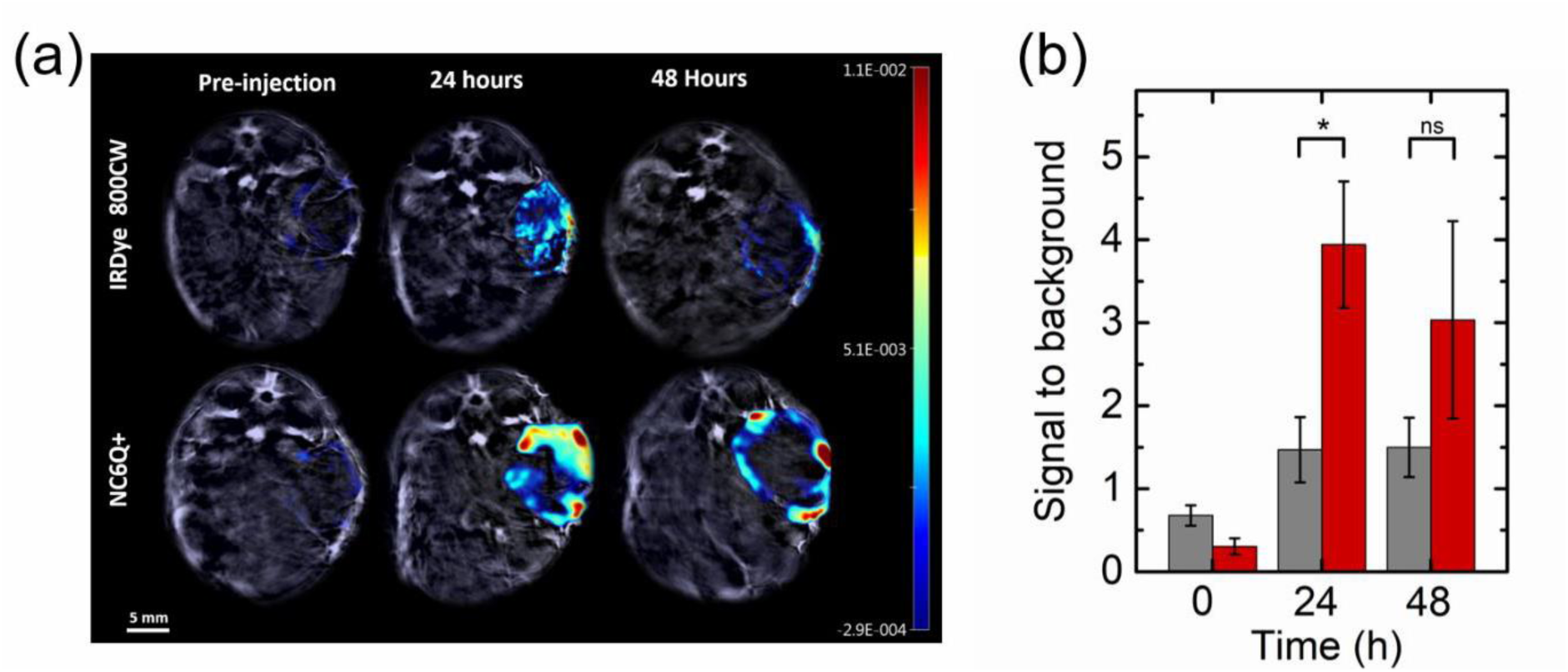
DNA NC show enhanced tumor uptake. (a) OT images with normalized mean pixel intensity (MPI) values obtained from mice before and at 24h and 48h after iv injection of either IRDye 800CW (top panel) or NC6Q+ (bottom panel). (b) Volumetric tumor-to-background ratio obtained from region of interest analysis of OT data before and at 24h and 48 hours after iv injection of either IRDye 800CW (grey data) or NC6Q+ (red data). Values are significantly different at 24h. n=3. *P<0.05. ns: not significant.

## 3. Discussion

Optoacoustic tomography (OT) enables the visualization of tumors with improved spatial resolution at greater depths than traditional optical imaging while maintaining advantages of non-invasiveness and low-cost. Molecular imaging is also possible with OT, upon administration of exogenous contrast agents, which are often based on NIR fluorophores derived from individual organic molecules.

Here, we demonstrated the potential of DNA-NCs to enhance the optoacoustic signal generation of small molecule dyes and prolong their clearance time, additionally enhancing their tumor accumulation. DNA-NCs possess unique characteristics to accomplish these properties afforded by their biocompatibility, as well as their fully customizable structure and size, with precise control for positioning of dye moieties. The reported DNA-NCs are able to provide a versatile platform to overcome the limitations of small molecule dyes, addressing both the low signal generation capability and rapid clearance.

Structural and optical characterization of DNA NCs showed excellent stability and optical response over time for the smaller NCs, including NC6Q+ and NC2Q+, with substantial enhancement of the optoacoustic signal generation. No impact on cellular proliferation rate was observed in the MDA-MB-231 cell line when incubated with the DNA NCs. Fast OT studies of heathy mice revealed the effect of the NC on the clearance rate. NC6Q+ exhibited a slower clearance rate when compared to NC2Q+ or the free dye, demonstrating the benefit of arranging free dyes into larger structures to prolong their circulation time in blood. The NC6Q+ structure displayed a 2.7-fold enhancement of OT signal when compared to the free fluorophore at the tumor site 24h after administration. Results at longer times (48 h) retained the trend but did not show a statistically significant improvement of the OT signal. It is interesting to note that compared to a previously reported DNA-origami structure for OT that used gold nanoparticles for signal generation,^[23]^ our approach exploits low molecular weight dyes for signal generation, which are already widely used in clinical trials. Furthermore, we utilized simpler DNA-based nanostructures, which affords the potential for scaling production to larger volumes should the approach become promising for clinical application.

While the present study has shown promise for the DNA-NC approach, there remain some limitations. The signal and circulation time enhancements remain relatively modest. The versatility of DNA nanotechnology permits the design of alternative NCs bearing a larger number of NIR dyes to further improve the optoacoustic enhancement provided by NC6Q+. To further enhance the circulation time, further size enlargement of the DNA-NC structure could be considered, yet in our hands we observed poorer stability of the larger construct size. In addition, while NC6Q+ is able to target the tumor, possibly aided by the EPR effect of solid tumors,^[42]^ further tailoring could be made in future to introduce specific moieties that enable a more selective targeting, to expand imaging capabilities towards other types of cancer and to enable a theranostic action upon integration of therapeutic agents.

## 4. Conclusions

In conclusion, we have shown that DNA nanotechnology can be used to generate nanocarriers that successfully enhance the OT signal by arranging pairs of IRDye 800CW fluorophores, thereby presenting a versatile and scalable approach for the fabrication of biocompatible contrast agents for deep tissue OT. Furthermore, we demonstrated the suitability of the presented nanocarrier NC6Q+ for *in vivo* OT in a breast cancer xenograft model. Hereby, NC6Q+ renders clear advantages over freely injected fluorophores by exhibiting an increased optoacoustic signal, a spectral response dependent on the H-dimer formation of IRDye 800CW, and successful accumulation at the tumor site delivering signal at 24 h, presenting enhanced OT contrast when compared to free fluorophores.

## 5. Experimental Section

### DNA NCs fabrication

Oligonucleotides were purchased from IDT (Integrated DNA Technologies, Inc.). Oligonucleotides modified with the IRDye 800CW were stored in IDTE buffer (Integrated DNA Technologies, Inc.), whereas oligonucleotides without modification were stored in RNAse-free water (Integrated DNA Technologies, Inc.). NC2, NC6 and NC8 were assembled at equimolar oligonucleotide concentrations in Phosphate Buffered Saline (PBS) solution. The assembly was carried out in a thermocycler using a thermal-annealing protocol adapted from previous reports^.[34]^ Namely, NC6 and NC8 were assembled by heating to 85° for 5 min, cooling down from 85 to 65°C in 20 steps (1°C per step, 5 min each step) and cooling down from 65 to 25°C in 80 steps (0.5°C per step, 12 min each step). NC2 was assembled under the following protocol:^[31, 32]^ 70 to 25 °C in 90 steps (0.5 °C per step, 30 s each step). The synthesized samples were stored at 4°C protected from light.

### DNA NCs structural characterization

The folding of the DNA NC was assessed by PAGE and DLS. For PAGE, DNA NCs were loaded and run for 80 minutes at 100 V in 10% polyacrylamide gel immersed in a solution containing 11 mM MgCl_2_ buffered with 0.5x TBE (pH=8.3). As a reference, a 50 bp ladder was run along with the samples. For subsequent visualization, the gels were stained in GelRed (Biotium, Fremont, CA, USA) and imaged under UV light transillumination. For the DLS measurements (Zeta Sizer, Malvern Instruments) the structures folded with unlabeled oligonucleotides (in order to avoid optical absorption of the sample) in PBS were measured at 1 μM of DNA NC concentration.

### DNA NCs stability in cell culture media

To establish the structural stability of NC6Q+ and NC8Q+ over time in cell culture media, the NCs were incubated in a solution of Dulbecco’s Modified Eagle’s medium (DMEM) supplemented with 10% of fetal calf serum (FCS). Incubation times were set to 0, 4, 8, 24 and 48hours. In all cases the incubation temperature was 37°C. Hereby, the NCs were incubated in DMEM+ 10% FCS to a final concentration of 250 nM for NC6Q+ and NC8Q+. The stability was determined using PAGE. The incubation mixtures were run for 80 minutes at 100V in a 10% polyacrylamide gel using 11 mM MgCl_2_ with 0.5x TBE (pH=8.3) as the running buffer. The degree of stability was expressed by the percentage of remaining full structure per lane in the gel using ImageJ. Specifically, for each incubation point the intensity of the band corresponding to the full structure was divided by the intensity of this band at time zero to yield the relative percentage.

### Optical absorbance and fluorescence measurements

Absorbance and fluorescence properties of the NCs were measured at 34°C. To compare the potential quenching effect of the NCQ+ versus NCQ-, the measurements were performed at matching IRDye 800CW concentration in PBS. Thus, the DNA NC concentration was prepared accordingly to match a final dye concentration of 2 µM (equivalent to DNA NC concentrations of 0.5 µM for NC6Q+, 1 µM for NC6Q-, 1 µM for NC2Q+ and 2 µM for NC2Q-). The absorbance measurements were carried out on a Varian Cary 300 Bio UV-vis Spectrophotometer and the fluorescence measurements were done with a Varian Cary Eclipse Fluorescence Spectrophotometer.

For the calculation of the quenching efficiency, the excitation was carried out at 778 nm. The percentage of quenching or quenching efficiency QE (%), was calculated at 795 nm of the emission spectra and averaged over three different samples for each type of NC as described by Equation 1, where IQ- and IQ+ are the intensities of the emission collected at the maximum of the fluorescence peak for the structures in the Q- and Q+ version, respectively.

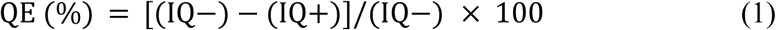

### Multispectral optoacoustic tomography in phantoms

A commercial multispectral OT system was employed (inVision256-TF; iThera Medical GmbH). The commercial OT system has been characterized and described previously.^[43]^ Briefly, for signal excitation the system uses a tunable (660–1300nm) optical parametric oscillator pumped by a nanosecond pulsed Nd:YAG laser operating at 10Hz repetition rate. Multi-element transducer array operating at 5 MHz center frequency, 60% bandwidth and with toroidal focusing is used as the detector. For phantom studies nanocarriers were encapsulated inside thin-walled optically transparent tubes located at 1 cm depth along the center of cylindrical tissue mimicking phantom. The tissue mimicking phantom was placed inside the temperature maintained (34°C) imaging chamber of the OT system. The phantoms were allowed to temperature stabilize for 10 minutes within the system prior to the initialization of the scan. OT data were acquired with 10 times frame averaging from multiple cross-sectional slices separated by a 1 mm step size to obtain averaged OT signal response over positions across the phantom. OT signals were acquired at 690, 705, 710, 719, 730, 760, 778, 800 and 850 nm excitation wavelengths. Mean pixel intensity (MPI) values were extracted from a region of interest (ROI) drawn within the thin walled plastic straw and the averaged values over the 5 scan positions at each wavelength. Averaged MPI values obtained at the different excitation wavelengths were plotted against the respective wavelengths to render multispectral optoacoustic spectra of the NCs. To determine the optoacoustic signal enhancement caused by H-dimer formation in the NCQ+ samples when compared to NCQ-, the area under the curve (AUC) for the obtained spectra were calculated to obtain the integrated optoacoustic intensity. The signal enhancement was calculated by using NCQ+ and NCQ-samples at specific DNA NC concentration to match the final dye concentration. Namely, 500 nM for NC6Q+, 1μM for NC6Q- were used to render a final dye concentration of 2 μM and 2 μM for NC2Q+,4 μM for NC2Q- were used to render a final dye concentration of 4 μM.

### Cell viability studies

MDA-MB-231 cells (passage 27) were grown in DMEM supplemented with 10% FCS and 50 U/ml of penicillin and 50 µg/ml of streptomycin. They were then seeded at a density of 60000 cells per ml of complete media into clear 48-well plates. DNA NCs were immediately added (4 wells per condition) at a final DNA concentration of 0.2 µM, 1 µM and 2 µM for NC2Q+ and 0.1 µM, 0.5 µM and 1 µM for NC6Q+. Wells were then imaged with an Incucyte Zoom live cells analysis system (Essen Bioscience). Nine phase contrast images per well were acquired every 3 hours and the cell confluence was measured for each time point using the Incucyte ZOOM confluence processing overview module (Essen Bioscience).

### Animal experiments

All animal procedures were conducted in accordance with the project (70-8214) and personal license (IFBB827BC) issued under the United Kingdom Animals (Scientific Procedures) Act, 1986. The procedures were reviewed by the Animal Welfare and Ethical Review Board at the CRUK Cambridge Institute under compliance form number CFSB1232V2. Immunodeficient female nude (BALB/c nu/nu) mice (Charles River) were used for conducting the biodistribution and tumor uptake studies. Tumor xenografts were inoculated subcutaneously on both flanks with 2.3 × 10^5^ cells (MDA-MB-231) in a final volume of 100?μL of 1:1 DMEM and matrigel (BD). Tumor growth was routinely monitored using calipers to ensure the tumor weight not exceeding 10% of body weight. For imaging and tail vein catheterization, mice were anaesthetized using <2.5% isoflurane in 100% oxygen, placed on a heat pad. For the administration of contrast agents, a catheter (made with 30G needle) was induced into the tail vein and was fixed in place using tissue glue (TS1050071F; TissueSeal). The mice were sacrificed by cervical dislocation towards the end of the study. Tumors and organs were immediately extracted for further histopathology investigations

### In vivo multispectral optoacoustic tomography and data quantification

For *in vivo* imaging, using the same commercial MSOT system as for phantom studies, anaesthetized animals were wrapped in a thin polyethylene membrane and placed in the animal holder, with thin water layer used as the acoustic coupling medium between skin and the polyethylene membrane. The holder was placed within the imaging chamber filled with degassed water which was maintained at 36 °C. The mice were allowed to temperature stabilize for 12 minutes within the imaging chamber of the OT system. The respiratory rate of the animals was maintained in the range 70-80 b.p.m. by modulating the isoflurane concentration (1.5-2.5% isoflurane concentration) as required during the entire scan. The animal holder was translated with a step size of 1 mm using a motorized stage to acquire transverse cross-sectional images. Multispectral optoacoustic images were acquired using 680, 700, 705, 715, 720, 730, 760, 777, 800, 825, 850 nm excitation wavelengths and 10 times frame averaging per wavelength. Each slice took 11?s to acquire, with overall imaging sessions lasting for a time ranging between 3 min to 1 hour.

Clearance of the NCs and free dye were assessed by performing fast OT of healthy female mice. Mice (n=3 for each group) were prepared according to the procedures described in the animal experiments section. NCs and free dye were administered via tail vein at consistent time points and injection rate with a dye concentration of 32 µM (equivalent to 16 µM of NC2Q+ and 8 µM of NC6Q+ for the structure). OT was continuously performed over 1 hour forming one cross-sectional image from the liver and kidney sections of the animal for approximately every 26 seconds. The signal contribution from the NCs was distinguished from the overall signal with linear unmixing, using the absorption spectra of the respective NCs, as well as with the optical absorption spectra of oxy-(HbO2) and deoxy-hemoglobin (Hb). Quantification of biodistribution and kinetics of the NCs over time were assessed by calculating the MPI obtained from ROI analysis of the right renal cortex and liver cross-sections of the animals. Kinetic profiles for the NCs and controls were smoothed using moving average with OriginPro 2016 software. Extracted metrics included: time to half maximum (T_1/2_) which gives the time taken for the signal to rise from baseline to 50% of maximum intensity; time to maximum (Tmax), time taken for the signal to rise from baseline to the maximum intensity; and time from maximum to half maximum (T_-1/2_), the time taken for the signal to decrease to 50% maximum intensity. Prior and subsequent to the MSOT measurements, fluorescence imaging of the animals was performed.

### Optoacoustic image and statistical analysis

OT images were reconstructed offline and analyzed using the ViewMSOT software package (v3.8; iThera Medical GmbH). The model-linear based reconstruction and linear regression based un-mixing (with pre-loaded spectra of the respective NCs, HbO_2_ and Hb values) within the software package were used. Images were reconstructed with a pixel size of 75µm x 75µm and 100µm × 100µm for the phantoms and mice respectively. The reconstructions accounted for the spatial and temporal impulse response of the transducer, with a pass band between 50 kHz and 7 MHz. Similar speed of sound values (as set by bringing the imaged object into focus using the wall of plastic tubes within phantoms or vessel within mice as targets) were used for all image reconstructions within a given data set.

For phantom studies, the region of interest (ROI) size and the positions used were identical across all data sets (see Figure S2 in Supporting Information). Mean optoacoustic signal intensity values given by the spectral components (NCs, HbO_2_ and Hb) were extracted from the manually drawn ROIs of the multispectral processed images. To assess the in vivo kinetics, ROIs were drawn around the right kidney cortex (to avoid any potential influence of light attenuation by the spleen) and the liver (see Figure S2c and d in supporting information) in each mouse. For tumor uptake studies, ROI’s that cover the entire tumor area were drawn on per slice basis to obtain the optoacoustic signal from the entire tumor volume (see Figure S2b in supporting information). Background ROIs were drawn on healthy tissue region near the spine for corresponding image slices. Signal-to-background (SBR) for the volume was calculated from averaging the ratio of the mean of the signal to the mean of the background values obtained from multiple cross-section images.

All statistical analyses were performed using GraphPad Prism (GraphPad Inc.). Uncertainty on mean values is represented by the standard error unless otherwise stated. Unpaired two tailed t -test was used for pairwise comparisons. P values of <0.05 were considered significant. *P<0.05, **P<0.01, ns: not significant.

## Acknowledgements

This work was funded by a Cancer Research UK Cambridge Centre Pump Prime Research Grant (KAZA/071 and KNZA-151). K. N. B. acknowledges funding by ERASMUS + and a PROMOS scholarship awarded by the DAAD. JJ and SEB are funded by the EPSRC-CRUK Cancer Imaging Centre in Cambridge and Manchester (C197/A16465); CRUK (C14303/A17197, C47594/A16267, C9545/A29580) and the EU-FP7-agreement FP7-PEOPLE-2013-CIG-630729. S. H. A. acknowledges funding by the University of Zaragoza (UZ2018-CIE-04); by Fundación Ibercaja y Universidad de Zaragoza (JIUZ-2018-CIE-04) and by the Gobierno de Aragón-FSE (E47_17R).

## Author Contributions

J.J., S.E.B. and S.H.A. proposed the concept and designed the experiments. K.N.B., A.P. and S.H.A. carried out the synthesis and characterization of nanostructures. J.J. and L.B. performed the cell culture studies. J.J. carried out the fluorescence and MSOT imaging in phantoms and *in vivo*. J. J., K. N. B., A. P., L. B. and S.H.A analyzed the data. J.J., K.N.B., S.E.B. and S.H.A. prepared and wrote the manuscript. All authors read the manuscript.

## Notes

### Competing Interest Statement

The authors have declared no competing interest.

## References

1. L. V. Wang, S. Hu, Science 2012, 335, 1458–1462.

2. A. Taruttis, V. Ntziachristos, Nat. Photonics 2015, 9, 219–227.

3. A. Taruttis, S. Morscher, N. C. Burton, D. Razansky, V. Ntziachristos, PLoS ONE 2012, 7, e30491.

4. X. L. Deán-Ben, D. Razansky, Light Sci. Appl. 2014, 3, e137.

5. D. Razansky, A. Buehler, V. Ntziachristos, Nat. Protoc. 2011, 6, 1121–1129.

6. C. Zhan, Y. Huang, G. Lin, S. Huang, F. Zeng, S. Wu, Small 2019, 15, 1900309.

7. V. Neuschmelting, S. Harmsen, N. Beziere, H. Lockau, H. T. Hsu, R. Huang, D. Razansky, V. Ntziachristos, M. F. Kircher, Small 2018, 14, 1800740.

8. S. K. Maji, S. Sreejith, J. Joseph, M. Lin, T. He, Y. Tong, H. Sun, S. W. K. Yu, Y. Zhao, Adv. Mater. 2014, 26, 5633–5638.

9. J. Weber, P. C. Beard, S. E. Bohndiek, Nat. Methods 2016, 13, 639–650.

10. M. Longmire, P. L. Choyke, H. Kobayashi, Nanomedicine 2008, 3, 703–717.

11. S. Roberts, M. Seeger, Y. Jiang, A. Mishra, F. Sigmund, A. Stelzl, A. Lauri, P. Symvoulidis, H. Rolbieski, M. Preller, J. Am. Chem. Soc. 2017, 140, 2718–2721.

12. J. Mérian, J. Gravier, F. Navarro, I. Texier, Molecules 2012, 17, 5564–5591.

13. S. M. Janib, A. S. Moses, J. A. MacKay, Adv. Drug Delivery Rev. 2010, 62, 1052–1063.

14. D. Peer, J. M. Karp, S. Hong, O. C. Farokhzad, R. Margalit, R. Langer, Nat. Nanotechnol. 2007, 2, 751.

15. J. Xie, S. Lee, X. Chen, Adv. Drug Delivery Rev. 2010, 62, 1064–1079.

16. Z. Yang, J. Song, W. Tang, W. Fan, Y. Dai, Z. Shen, L. Lin, S. Cheng, Y. Liu, G. Niu, P. Rong, W. Wang, X. Chen, Theranostics 2019, 9, 526–536.

17. S. Sreejith, J. Joseph, M. Lin, N. V. Menon, P. Borah, H. J. Ng, Y. X. Loong, Y. Kang, S. W.-K. Yu, Y. Zhao, ACS nano 2015, 9, 5695–5704.

18. K. T. Nguyen, S. Sreejith, J. Joseph, T. He, P. Borah, E. Y. Guan, S. W. Lye, H. Sun, Y. Zhao, Part Part Syst Char. 2014, 31, 1060–1066.

19. Y. Matsumura, H. Maeda, Cancer Res. 1986, 46, 6387–6392.

20. T. Stylianopoulos, Ther. Deliv. 2013, 4, 421–423.

21. A. V. Pinheiro, D. Han, W. M. Shih, H. Yan, Nat. Nanotechnol. 2011, 6, 763–772.

22. D Mathur, I. Medintz, Adv. Healthcare Mater. 2019, 8, 1801546.

23. Y. Du, Q. Jiang, N. Beziere, L. Song, Q. Zhang, D. Peng, C. Chi, X. Yang, H. Guo, G. Diot, V. Ntziachristos, B. Ding, J. Tian, Adv. Mater. 2016, 28, 10000–10007.

24. Q. Hu, H. Li, L. Wang, H. Gu, C. Fan, Chem. Rev. 2018, 119, 6459–6506.

25. V. Kumar, S. Palazzolo, S. Bayda, G. Corona, G. Toffoli, F. Rizzolio, Theranostics 2016, 6, 710–725.

26. Q. Jiang, S. Zhao, J. Liu, Linlin Song, Zhen-Gang Wang, B. Ding, Adv. Drug Deliv. Rev. 2019, 141, 2–21.

27. Q. Zhang, Q. Jiang, N. Li, L. Dai, Q. Liu, L. Song, J. Wang, Y. Li, J. Tian, B. Ding, ACS nano 2014, 8, 6633–6643.

28. K. R. Kim, Y. D. Lee, T. Lee, B. S. Kim, S. Kim, D. R. Ahn, Biomaterials 2013, 34, 5226-5235.

29. D. Jiang, Y. Sun, J. Li, Q. Li, M. Lv, B. Zhu, T. Tian, D. Cheng, J. Xia, L. Zhang, L. Wang, Q. Huang, J. Shi, C. Fan, ACS Appl. Mater. Interfaces 2016, 8, 4378–4384.

30. D. Jiang, Z. Ge, H.-J. Im, C. G. England, D. Ni, J. Hou, L. Zhang, C. J. Kutyreff, Y. Yan, Y. Liu, S. Y. Cho, J. W. Engle, J. Shi, P. Huang, C. Fan, H. Yan, W. Cai, Nat. Biomed. Eng. 2018, 2, 865–877.

31. K. N. Baumann, A. C. Fux, J. Joseph, S. E. Bohndiek, S. Hernández-Ainsa, Chem. Commun. 2018, 54, 10176–10178.

32. J. Joseph, K. N. Baumann, P. Koehler, T. J. Zuehlsdorff, D. J. Cole, J. Weber, S.. E. Bohndiek, S. Hernández-Ainsa, Nanoscale 2017, 9, 16193–16199.

33. M. Laramie, M. Smith, F. Marmarchi, L. McNally, M. Henary, Molecules 2018, 23, 2766.

34. K. Göpfrich, T. Zettl, A. E. C. Meijering, S. Hernández-Ainsa, S. Kocabey, T. Liedl, U. F. Keyser, Nano Letters 2015, 15, 3134–3138.

35. B. Wei, M. Dai, P. Yin, Nature 2012, 485, 623–626.

36. H. Kobayashi, M. R. Longmire, M. Ogawa, P. L. Choyke, Chem. Soc. Rev. 2011, 40, 4626–4648.

37. M. Ogawa, N. Kosaka, C. A. S. Regino, M. Mitsunaga, P. L. Choyke, H. Kobayashi, Mol. BioSyst. 2010, 6, 888–893

38. M. Ogawa, N. Kosaka, P. L. Choyke, H. Kobayashi, ACS Chem. Biol. 2009, 4, 535–546.

39. A. Ohmann, C.-Y. Li, C. Maffeo, K. Al Nahas, K. N. Baumann, K. Göpfrich, J. Yoo, U. F. Keyser, A. Aksimentiev, Nat. Commun. 2018, 9, 2426.

40. S. M. Moghimi, A. C. Hunter, J. C. Murray, Pharmacol Rev. 2001, 53, 283–318.

41. J. L. Kovar, W. Volcheck, E. Sevick-Muraca, M. A. Simpson, D. M. Olive, Anal. Biochem. 2009, 384, 254–262.

42. J. Fang, H. Nakamura, H. Maeda, Adv. Drug Delivery Rev. 2011, 63, 136–151.

43. J. Joseph, M. R. Tomaszewski, I. Quiros-Gonzalez, J. Weber, J. Brunker, S. E. Bohndiek, J. Nucl. Med. 2017, 58, 807.

